# Predicting Antimicrobial Resistance Using Conserved Genes

**DOI:** 10.1101/2020.04.29.068254

**Authors:** Marcus Nguyen, Robert Olson, Maulik Shukla, Margo VanOeffelen, James J. Davis

## Abstract

A growing number of studies have shown that machine learning algorithms can be used to accurately predict antimicrobial resistance (AMR) phenotypes from bacterial sequence data. In these studies, models are typically trained using input features derived from comprehensive sets of known AMR genes or whole genome sequences. However, it can be difficult to determine whether genomes and their corresponding sets of AMR genes are complete when sequencing contaminated or metagenomic samples. In this study, we explore the possibility of using incomplete genome sequence data to predict AMR phenotypes. Machine learning models were built from randomly-selected sets of core genes that are held in common among the members of a species, and the AMR-conferring genes were removed based on their protein annotations. For *Klebsiella pneumoniae*, *Mycobacterium tuberculosis*, *Salmonella enterica*, and *Staphylococcus aureus*, we report that it is possible to classify susceptible and resistant phenotypes with average F1 scores ranging from 0.80-0.89 with as few as 100 conserved non-AMR genes, with very major error rates ranging from 0.11-0.23 and major error rates ranging from 0.10-0.20. Models built from core genes have predictive power in the cases where the primary AMR mechanism results from SNPs or horizontal gene transfer. By randomly sampling non-overlapping sets of core genes for use in these models, we show that F1 scores and error rates are stable and have little variance between replicates. Potential biases from strain-specific SNPs, phylogenetic sampling, and imbalances in the phylogenetic distribution of susceptible and resistant strains do not appear to have an impact on this result. Although these small core gene models have lower accuracies and higher error rates than models built from the corresponding assembled genomes, the results suggest that sufficient variation exists in the core non-AMR genes of a species for predicting AMR phenotypes. Overall this study suggests that building models from conserved genes may be a potentially useful strategy for predicting AMR phenotypes when genomes are incomplete.

## Introduction

The discovery and use of antimicrobial agents for the treatment of bacterial infections revolutionized medicine in the twentieth century. In 1900, the top three causes of death in the United States were pneumonia, tuberculosis, and diarrhea/enteritis^1^. Antimicrobial therapy, coupled with medical advancements and sanitation improvements, has resulted in a marked shift in these statistics, with the top three causes of death in the United States now being heart disease, cancer, and unintentional injuries^2,3^. The rise of antimicrobial resistance (AMR), along with the sluggish development of new antimicrobial drugs, threatens to jeopardize this achievement.

Antimicrobial susceptibility testing is the gold standard for determining which antibiotics will be effective against bacterial pathogens. This requires culturing the organisms in the presence of a panel of antibiotics^4,5^. Since culturing can be slow, clinicians often rely on empirical judgement when administering antibiotics. When the administration of antibiotics is incorrect or inappropriate, it can increase mortality rates and exacerbate the spread of AMR^6,7^. Developing diagnostics that can determine the AMR phenotypes of bacterial pathogens in real time is crucial for reducing morbidity and mortality in patients, and for lowering endemic levels of AMR through more precise antibiotic prescription and stewardship practices^8,9^.

With the continued reductions in cost and the development of devices that are more suitable for point of care use, genome sequencing has received attention for its potential value as a diagnostic tool^10–12^. Many bioinformatic techniques have been developed for making sequence-based comparisons against large comprehensive databases of known AMR genes, proteins, and variants making the prediction of resistant phenotypes possible^13–15^. These predictions are typically made using either rules-based or machine learning models^16–20^. Several studies have also built machine learning models for predicting AMR phenotypes by using assembled genomes or pan genomes as training sets^21–27^. In these cases, the machine learning algorithm detects the most discriminating features (typically short nucleotide k-mers) from a training set with laboratory-derived AMR phenotypes. Both the AMR gene- and whole genome-based approaches have the limitation that they require either a complete genome or the complete set of AMR genes from a genome to provide accurate AMR phenotype predictions.

Although whole genome sequencing of bacterial isolates provides extensive information about AMR, pathogenicity, and epidemiology, it also requires a culturing step, which means that it is not much faster than conventional susceptibility testing. Thus, culture-free diagnostic techniques, such as shotgun metagenomics and PCR-based amplification of AMR markers, represent appealing alternatives, but these also come with challenges. For instance, in metagenomics, where the pathogen DNA is sequenced directly from an infection source, there can be difficulty in eliminating contaminating host DNA, accurately binning reads or contigs into individual genomes, determining if binned genomes are complete, and assessing the risk posed by incomplete genomes found in the sample. Furthermore, while whole genome sequencing of pure cultures enables accurate source attribution for mobile genetic elements carrying AMR genes, this becomes more difficult in a metagenomic sample^28^. PCR-based approaches, including the direct amplification of AMR genes, or amplification paired with sequencing, have similar challenges including the difficulties in amplifying a comprehensive set of AMR genes or regions, and the subsequent ability to attribute these detected AMR genes to specific pathogens. Given the obvious appeal and drawbacks of culture-free diagnostic approaches, developing strategies for predicting AMR phenotypes from incomplete or mixed genome sequence data presents an interesting technical challenge.

In a recent study, we used the complete assembled genomes of 1667 *Klebsiella pneumoniae* clinical isolates to build machine learning models for predicting AMR phenotypes for 20 antibiotics^21^. During this study, we did an experiment where we built similar model from the same set of genomes, except that we excluded the known AMR genes based on their annotations. To our surprise, the resulting model had nearly identical accuracies and error rates across all antibiotics compared with the model built from the full genomes (approximately 92%). This suggests that it may be possible to build accurate models with partial genome sequence data. In this study, we explore this problem in greater detail.

## Materials and Methods

### Data acquisition

Whole genome sequence data and paired laboratory-derived antimicrobial susceptibility test (AST) data were downloaded from the PATRIC FTP server (ftp://ftp.patricbrc.org/RELEASE_NOTES/PATRIC_genomes_AMR.txt) on or after December 1, 2018^29,30^. We chose to analyze *Klebsiella pneumoniae*, *Mycobacterium tuberculosis*, *Salmonella enterica*, and *Staphylococcus aureus*, which had the largest collections of genomes paired with AST data in this collection at the time (Table 1, Tables S1-6). The corresponding genomic data were also downloaded from the PATRIC FTP server (ftp://ftp.patricbrc.org/genomes)^31^. The PATRIC command line interface was used for all other annotation and metadata acquisition tasks^32^. In the case of *M. tuberculosis*, we also used data from a study that had not yet been integrated into the PATRIC collection^33^. The reads for these genomes were downloaded from the European Nucleotide Archive^34^, assembled using the full spades pipeline at PATRIC^32,35,36^, and annotated using the PATRIC annotation service, which is a variant of RAST^37^.

**Table 1.**
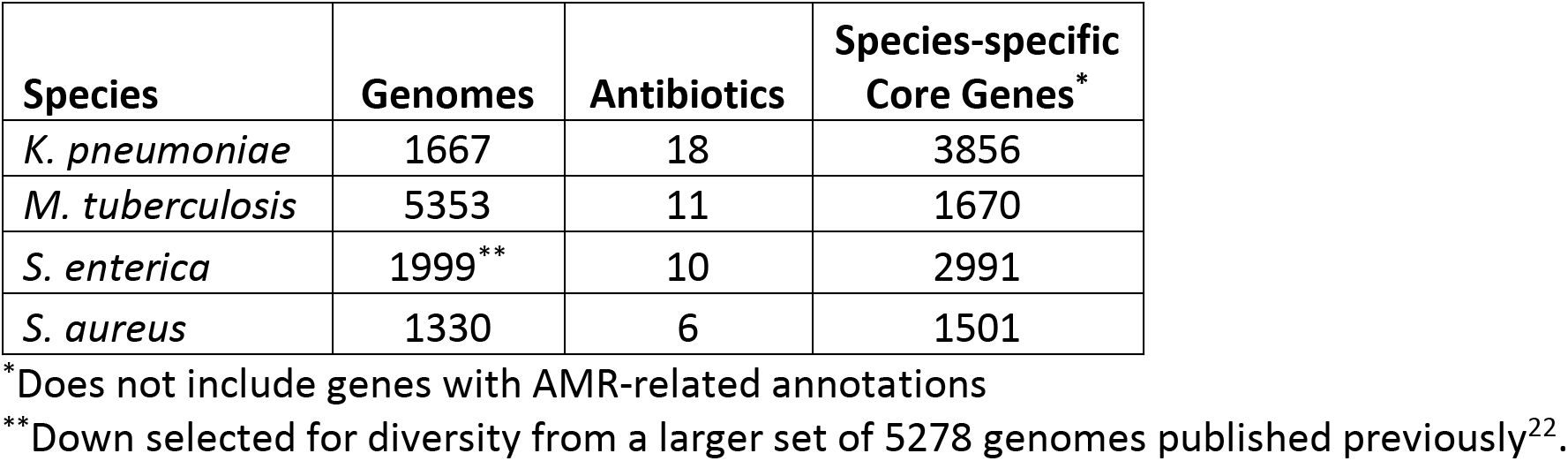
Data sets used in this study.

AMR phenotype data in the PATRIC collection usually exists in two forms, the first being a laboratory derived value such as a minimum inhibitory concentration (MIC), and the second being the susceptible, intermediate, or resistant determinations that were made by the authors at the time of the study. When only the susceptible or resistant (SR) determination was available for a genome, we did not mix data based on breakpoint values from the Clinical and Laboratory Standards Institute (CLSI) and the European Committee on Antimicrobial Susceptibility Testing (EUCAST) because their recommended breakpoint values can differ^38,39^. When AMR phenotypes were published as MICs, they were converted to SR determinations using either CLSI or EUCAST guidelines to create the largest possible set of genomes with SR phenotypes^38,39^. The use of CLSI or EUCAST breakpoints for a given dataset was determined on the total size of the dataset, with the larger dataset being chosen. Classification was not performed on intermediate phenotypes because they are underrepresented. All SR data used in this study are shown in Tables S1-4, and MICs are shown in Tables S5 and S6.

### Core conserved gene sets

In order to work with clean subsets of genomes, we chose to base analyses on the protein-encoding genes that are shared among members of the same species. We used the “PATtyFam” collection, which is a set of protein families that cover the ~230,000 publicly available genomes in the PATRIC database^40^. Protein similarity for building these families is based on the RAST signature k-mer collection^37^, and all proteins must have the same annotation in order to be members of the same family. Specifically, we used the genus-specific PATRIC local families (PLFs). All genomes were reannotated so that they had the same set of protein family calls based on the May 31, 2018 protein family build, using the PATRIC annotation server^37^. All families with a PATRIC annotation associated with AMR were removed from consideration in this study^30^.

We used two criteria when defining core gene sets. First, for each family, the average nucleotide length was computed for the corresponding genes. Any family member that had a total nucleotide length that was less than half of the average length, or that was 50% longer than the average length, was excluded. This helped to eliminate duplicate genes, partial genes, and mixtures of genes encoding single and multi-subunit proteins. Next, we excluded any family whose members represented less than 99% of the genomes of the set. This criterion was relaxed to 75% to build sufficient core family sets for *S. aureus.* After the core gene sets were computed, sets of 25, 50, 100, 250, and 500 core genes were randomly selected for each model.

### Model generation

K-mer-based models were generated using XGBoost (XGB)^41^ to classify susceptible and resistant phenotypes, or to predict MICs following protocols described previously^21,22^. Briefly, the sequences of core genes or whole assembled genomes were converted into nucleotide k-mers, with a step size of one nucleotide, and lexicographically sorted and counted using the k-mer counting program KMC^42^. Partial k-mers at the ends of sequences, and sequences containing ambiguous nucleotides were not considered. 15-mers were used to compute models for core genes, and 10-mers were used to build models from assembled genomes to ensure that all models would fit in memory^22^. For the *S. enterica* models, a diverse set of 1999 genomes was chosen for analysis out of an original set of 5278 genomes as described previously^22^, so that models could be computed efficiently.

Matrix files were constructed by merging the k-mer counts and the one-hot encoded phenotype and antibiotics. That is, each row of the matrix contains all of the k-mer counts for a given genome’s core genes, the encoded antibiotic, and the phenotype for a single genome-antibiotic pair. Because 10-mers are more likely to be redundant in a genome than 15-mers, we use k-mer counts when building 10-mer based models and presence versus absence of a given k-mer when building 15-mer-based models.

We have shown previously that in this context, the XGB model parameter that has the greatest effect on model accuracy is tree depth^21^. This parameter was varied to tune the models. Unless otherwise stated, core gene models reported used a depth of sixteen, and whole genome models were tuned to a depth of four. The learning rate was set to 0.0625, and column and row subsampling was set to 1.0 as described previously^21,22^. The number of rounds of boosting was limited to 1000. Since susceptible and resistant classes are not balanced, we weighted the counts for each class using the formula, 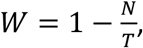 where *N* is count of a given class for an antibiotic, and *T* is the total counts for all classes for the given antibiotic. This weighting was not used for the MIC models or tree-based analyses described below.

Models were evaluated by dividing the set of genomes for each species into non-overlapping training (80%), testing (10%), and validation (10%) sets. Unless otherwise stated, statistics including the accuracy and macro-averaged F1 scores are shown for the first five folds of a 10-fold cross validation with 95% confidence intervals (CI). This was done to reduce the large compute time over the many sample replicates and parameter optimizations depicted in this study. To prevent overfitting, the accuracy of the model on the held-out validation set was monitored and training was stopped if the validation set accuracy did not improve after 25 rounds of boosting. Models were assessed by computing the accuracies, 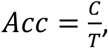 where *C* is the correctly predicted samples and *T* is the total samples tested, or the macro-averaged F1 score, 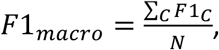 where *C* is a class and *N* is the number of classes. The very major error rate (VME), which is the fraction of resistant test set genomes that is predicted to be susceptible, and the major error rate (ME), which is the fraction of susceptible test set genomes that are predicted to be resistant, were also used to evaluate models^43^. For MIC models, accuracy is depicted within ± 1 two-fold dilution step, which is the limit of resolution for most automated laboratory AST devices^43^.

To control for unforeseen effects relating to the methodology or algorithm choice, we also built models from nucleotide alignments following a similar protocols to those that have been described previously^25,26^. One random subset of 100 core genes per species was chosen for the analysis. Each of the 100 core genes sets was aligned using MAFFT version 7.13^44^. Alignment columns where ≥95% of the characters were dashes (i.e., gaps) were masked. Start positions were trimmed to the first column that had no dashes in order to eliminate variability from start site calling, rather than true nucleotide differences. Any genome missing one or more of the 100 core genes was removed. Columns with 100% conservation were eliminated because they lack discriminating information. Alignments were then concatenated and one-hot encoded so that each nucleotide, the N character, and the dash character were encoded in a 6-digit string. The one-hot encoded alignments and AST phenotypes were then merged to form a matrix. XGB and Random Forest (RF)^45^ were then used to build the classifiers. For XGB, the same model parameters were used as described above for the k-mer-based core gene models. RF parameters were set to bagging of 1000 decision trees each, at a depth of 16, subsampling 75% of rows, and subsampling 75% of features. Accuracies and error rates were generated using cross validations as described above. To assess the impact that strain-specific SNPs had on these models, alignment columns with varying fractions of conservation were incrementally removed, and the models were recomputed with accuracy statistics. The Nucleotide alignments, one-hot encoded alignments, and matrix files are available on our GitHub page: https://github.com/jimdavis1/Core-Gene-AMR-Models.

### Tree-based analyses

Using too many closely related strains, or having closely related strains with skewed S or R phenotype distributions, could result in biased models. To assess this, we generated phylogenetic trees for each of the four species and computed models that were weighted based on the total number of tips in a subtree, and the distributions of S and R classes within each subtree. By defining subtrees at different tree depths, potential biases can be assessed in more closely or distantly related sets of organisms.

Trees were generated for each species by randomly selecting a set of 100 leftover core genes that were not used for computing the core gene models. Concatenated nucleotide alignments were generated as described above, and computed with FastTree using the nucleotide and generalized time reversible model options^46^. The trees were then divided into all possible subtrees in order to define clades of related genomes of various sizes. Each subtree containing more than one tip was midpoint rooted, and the tree distance was used to define the size of the subtree. At a small distance, subtrees will be comprised of nearly identical tips. As the distance increases, subtree membership becomes more inclusive, with more diverse sequences being represented within a subtree, until ultimately the distance becomes so large that all the tips in the tree are represented by a single subtree. In this way, by defining the most inclusive subtrees at increasing tree distances, we can measure the effect of imbalances in diversity and SR distribution on the models at varying levels of phylogenetic resolution.

K-mer based XGB models from core genes were created as described above. To weight genomes based on subtree size, the number of genomes in the subtree was divided by the total number of genomes in the model, and the corresponding fraction was assigned to each genome as a weight, using the equation, 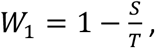 where *S* is the size of the subtree a genome belonged to and *T* is the total number of genomes analyzed. Likewise, the fraction of susceptible and resistant genomes was computed for each subtree, and a value of 1 minus the appropriate fraction was assigned to each genome as a weight, using the equation, 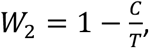 where C is the number of genomes of the same class within the subtree as the genome being weighted and T is the total number of genomes within a given subtree. However, if *C* = *T*, then the weight assigned was 1 (no weighting). In this way, frequently occurring phenotypes within a clade had low weights, and rarely occurring phenotypes had high weights. Finally, to assess both features together, the subtree size weight was multiplied by the SR weight.

### Analysis of top features

Ten k-mer-based XGB models were built from 100 randomly-selected non-overlapping core genes as described above. The k-mers that were used by each model, and their associated feature importance values were obtained. For each gene in the model, the corresponding feature importance of each corresponding k-mer was summed to generate a total feature importance for the gene. The location of each gene was plotted using its coordinates in a high-quality reference genome.

## Results

### AMR models based on core genes have predictive power

In previous work, we observed that is possible to build accurate AMR phenotype prediction models from whole genomes without using the AMR genes^21^. In this study, in order to explore the possibility of building models from limited genome sequence data, we chose to build models from core genes that are held in common among the members of a species, and which are not annotated as having a direct role in AMR^37,40^. By being nearly universally conserved, core genes are less likely to be horizontally transferred, and are also useful for assessing genome completeness and phylogeny. We built machine learning models using the core gene sets for *K. pneumoniae, M. tuberculosis, S. enterica,* and *S. aureus,* which have a large number of publicly available genomes with laboratory-derived AMR metadata (Tables 1-2, S1-6). For all species, classifiers were built for predicting susceptible and resistant (SR) phenotypes. We used the XGBoost (XGB)^41^ machine learning algorithm as described previously and 15-mer oligonucleotide k-mers from the core gene sets along with the SR phenotypes as features to train each model^21,22^.

**Table 2.** Counts of susceptible and resistant genomes used in this study, data are displayed as (Susceptible|Resistant).

| Antibiotic | Abv. | K.<br>pneumoniae | M.<br>tuberculosis | S.<br>enterica | S. aureus |
| --- | --- | --- | --- | --- | --- |
| Amikacin | AMK | 1320 \| 103 | 868 \| 230 |  |  |
| Amoxicillin/Clavulanate | AMC |  |  | 1489 \| 400 |  |
| Ampicillin | AMP |  |  | 1291 \| 706 |  |
| Ampicillin/Sulbactam | SAM | 90 \| 1455 |  |  |  |
| Aztreonam | ATM | 216 \| 1407 |  |  |  |
| Capreomycin | CAP |  | 846 \| 214 |  |  |
| Cefazolin | CFZ | 97 \| 1570 |  |  |  |
| Cefepime | FEP | 418 \| 963 |  |  |  |
| Cefoxitin | FOX | 667 \| 828 |  | 1599 \| 348 |  |
| Ceftazidime | CAZ | 136 \| 1488 |  |  |  |
| Ceftiofur | TIO |  |  | 1602 \| 393 |  |
| Ceftriaxone | CRO | 80 \| 1528 |  | 1601 \| 397 |  |
| Cefuroxime sodium | CXM | 91 \| 1469 |  |  |  |
| Chloramphenicol | CHL |  |  | 1864 \| 88 |  |
| Ciprofloxacin | CIP | 201 \| 1424 |  |  | 752 \| 243 |
| Clindamycin | CLI |  |  |  | 444 \| 144 |
| Erythromycin | ERY |  |  |  | 1016 \| 257 |
| Ethambutol | EMB |  | 3889 \| 484 |  |  |
| Ethionamide | ETO |  | 167 \| 123 |  |  |
| Fusidic acid | FA |  |  |  | 1156 \| 83 |
| Gentamicin | GEN | 926 \| 683 |  | 1639 \| 329 |  |
| Imipenem | IPM | 1160 \| 478 |  |  |  |
| Isoniazid | INH |  | 4098 \| 1204 |  |  |
| Kanamycin | KAN |  | 614 \| 167 |  |  |
| Levofloxacin | LVX | 349 \| 1287 |  |  |  |
| Meropenem | MEM | 1134 \| 481 |  |  |  |
| Methicillin | MET |  |  |  | 771 \| 215 |
| Moxifloxacin | MXF |  | 593 \| 85 |  |  |
| Ofloxacin | OFX |  | 182 \| 176 |  |  |
| Penicillin | PEN |  |  |  | 175 \| 1063 |
| Piperacillin/Tazobactam | TZP | 432 \| 1048 |  |  |  |
| Pyrazinamide | PZA |  | 3605 \| 428 |  |  |
| Rifampin | RIF |  | 4438 \| 828 |  |  |
| Streptomycin | STR |  | 2140 \| 684 | 381 \| 772 |  |
| Sulfisoxazole | FIS |  |  | 1108 \| 772 |  |
| Tetracycline | TET | 739 \| 778 |  | 867 \| 1124 |  |
| Tobramycin | TOB | 589 \| 723 |  |  |  |
| Trimethoprim/Sulfamethoxazole | SXT | 416 \| 1251 |  |  |  |

For each species, we started by randomly selecting subsets of core genes ranging in size from 25-500 genes. We then built SR classifiers for each set, tuning the XGB parameter for tree depth, which has been shown previously to have the most influence on models of this type^21^ (Figure S1). A tree depth of 16 was chosen for the models because the F1 scores tend to plateau beyond this point. In most cases, we see little improvement beyond depths of 16 regardless of gene set size, so it is likely that we are nearing the maximum accuracy that these gene sets can provide given the genes in the sets, set sizes, and experimental design. For all four species, models built from 25 genes and optimized to a tree depth of 16, range in their average F1 scores from 0.75 [0.73-0.77, 95%CI] for *S. enterica* to 0.80 [0.78-0.81, 95%CI] for *K. pneumoniae* (Figure 1). The F1 scores increase as the set size increases, with the models built from 500 genes having F1 scores ranging from 0.84 [0.81-0.86, 95%CI] for *M. tuberculosis* to 0.89 [0.86-0.90, 95%CI] for *S. aureus.* The average very major error rate (VME), which is defined as resistant genomes that are erroneously predicted to be susceptible, and the average major error rate (ME), which is defined as susceptible genomes that are erroneously predicted to be resistant, tend to go down as gene set size increases. Although the core gene set models described in Figure 1 have lower F1 scores and higher error rates than full-genome models that have been published previously^21–24,27,29^, their accuracies are striking given the small sizes of the input data sets and the removal of well-annotated AMR genes.

**Figure 1.**
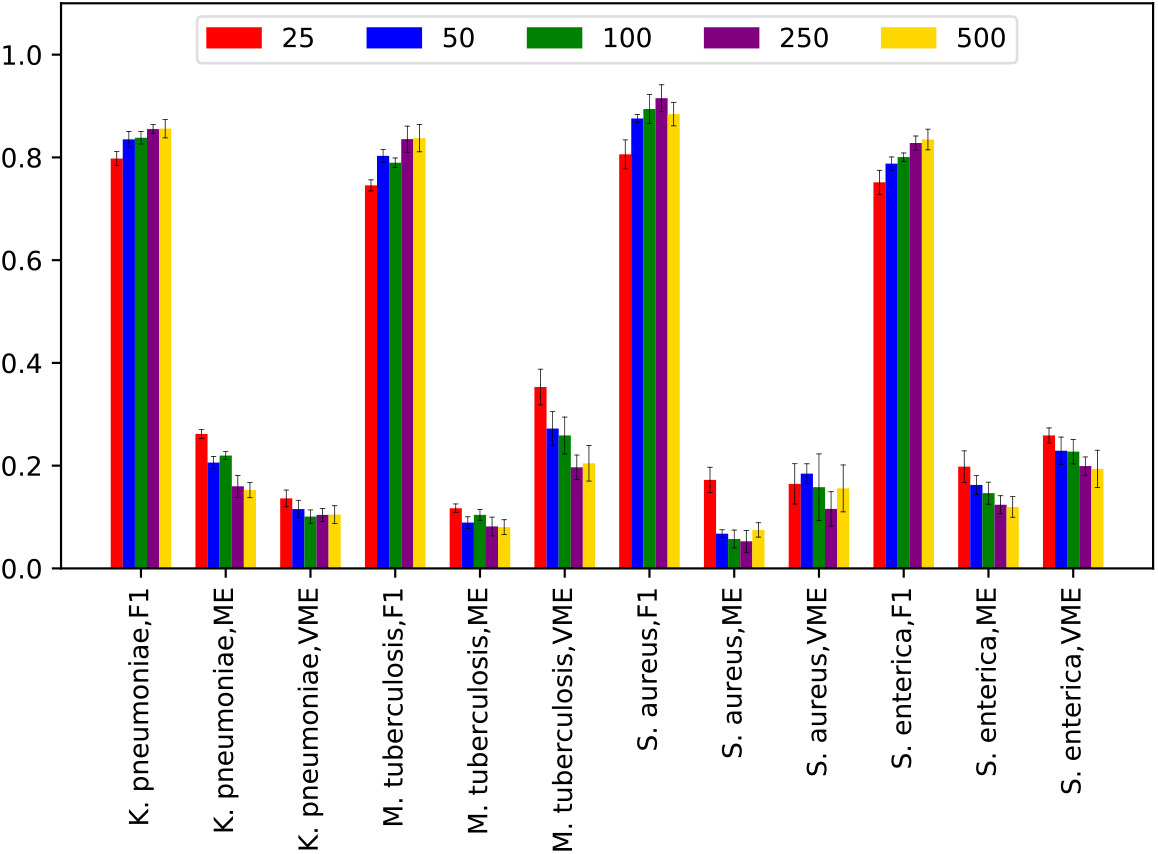
F1 scores, very major error (VME), and major error (ME) rates for AMR phenotype classifiers built from core gene sets. Randomly selected core gene sets ranging in size from 25-500 genes are shown. Error bars depict 95% confidence intervals.

### Models built from randomly sampled core gene sets are consistent

To assess the variability that could be expected from building models from core gene sets, we built models by randomly selecting non-overlapping sets of genes. Ten models, each containing 100 non-overlapping core genes, were computed for each species and the accuracies, F1 scores, and error rates were averaged over all 10 models (Figure 2, Table S7). The average F1 scores for these 100-gene models range from 0.80 in *M. tuberculosis* and *S. enterica* to 0.89 for *S. aureus*, and are 5-17% lower than the accuracies for models built from the same set of genomes using the whole assembled genomes as input. Within a species, we observe little variation in the accuracies or F1 scores for each gene set. In each of the four species, the average 95% confidence intervals for each model differ by only 1-2% for all ten replicates. This indicates that most randomly-selected subsets of 100 core genes will have accuracies within this range, regardless of the functions encoded in the underlying gene sequences.

**Figure 2.**
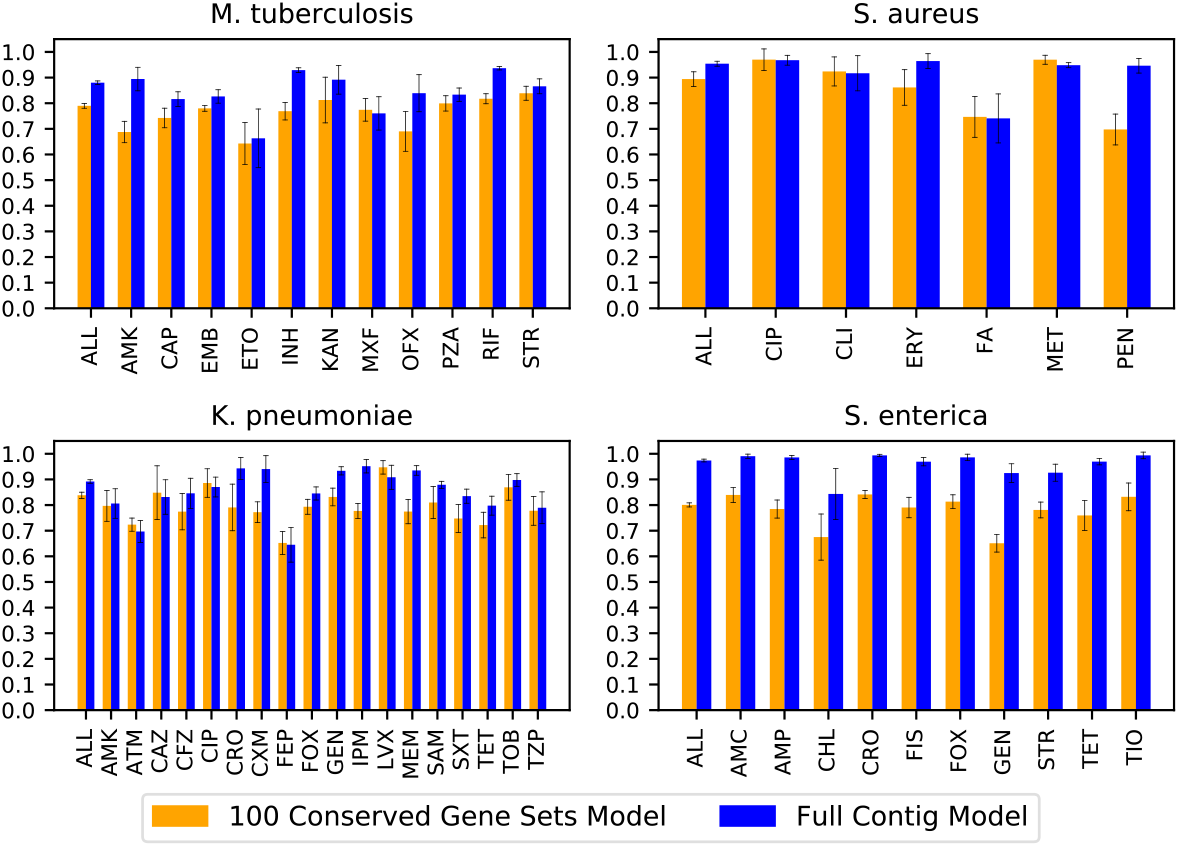
Average F1 scores by antibiotic for ten models built from non-overlapping sets of 100 core genes (orange bars) compared with the F1 scores for models built from whole assembled contigs for the same genomes (blue bars). Error bars represent 95% confidence intervals for the whole genome model, and the minimum and maximum observed in all ten replicates for the core gene models.

Although the F1 scores are higher for models built from whole genomes, the models built from core genes show a similar pattern across antibiotics. That is, when a F1 score for an antibiotic is high in a whole genome model, it also tends to be high in the 100-gene set models and vice versa (Figure 2). There are also many examples where the accuracy is high regardless of whether the AMR mechanism is known to result from chromosomally-encoded SNPs in core genes, or horizontal gene transfer. For instance, the 100 core gene models have average F1 scores of 0.90 [0.88-0.91, 95%CI] and 0.96 [0.96-0.97, 95%CI] for predicting ciprofloxacin resistance in *K. pneumoniae* and *S. aureus*, respectively. Ciprofloxacin resistance is often caused by SNPs in the *gyrA* gene^47^, but this gene is not used in any of the ten subsamples for either species. This means that other core genes carry sufficient information for making the prediction. This is also true in cases where the AMR mechanism is the result of horizontal gene transfer. One notable example is methicillin resistance in *S. aureus*, where resistance genes are carried by SCC*mec* elements^48^, which had an F1 score of 0.96 [0.95-0.97, 95% CI]. Another example is cephalosporin resistance in *S. enterica*, which has F1 scores ranging from 0.82-0.84, and is known to be plasmid-mediated^49^ (Table S7). The annotated functions for each of the ten core gene subsets are listed in Table S8 for each species.

In nearly all cases, the error rates are higher for the core gene models than they are for the whole genome models. The average VME rates for the whole genome models range from 0.04 [0.03-0.05, 95%CI] in *S. enterica* to 0.12 [0.11-0.12, 95%CI] for *M. tuberculosis,* and in the core gene models, they range from 0.11 [0.10-0.12, 95%CI] in *K. pneumoniae* to 0.23 [0.21-0.24, 95%CI]for *M. tuberculosis* (Figure 3, Table S7). Likewise, the ME rates are mostly higher in the core gene models than the whole genome models (Figure 4). The average ME rates for the whole genome models range from 0.01 [0.01-0.02, 95%CI] for *S. enterica* to 0.12 [0.11-0.13, 95%CI] for *K. pneumoniae.* In the 100 gene models, the average ME rates range from 0.06 [0.05-0.07, 95%CI] in *S. aureus* to 0.20 [0.18-0.22, 95%CI] in *K. pneumoniae* (Table S7, Figure 4). Overall, the antibiotics with poor F1 scores also tend to have higher error rates. There are also larger confidence intervals, and thus more variability predicting S or R phenotypes for antibiotics with an underrepresented class. Based on Figure 1, we would expect the VME and ME rates to go down as the core gene set size is increased beyond 100 genes.

**Figure 3.**
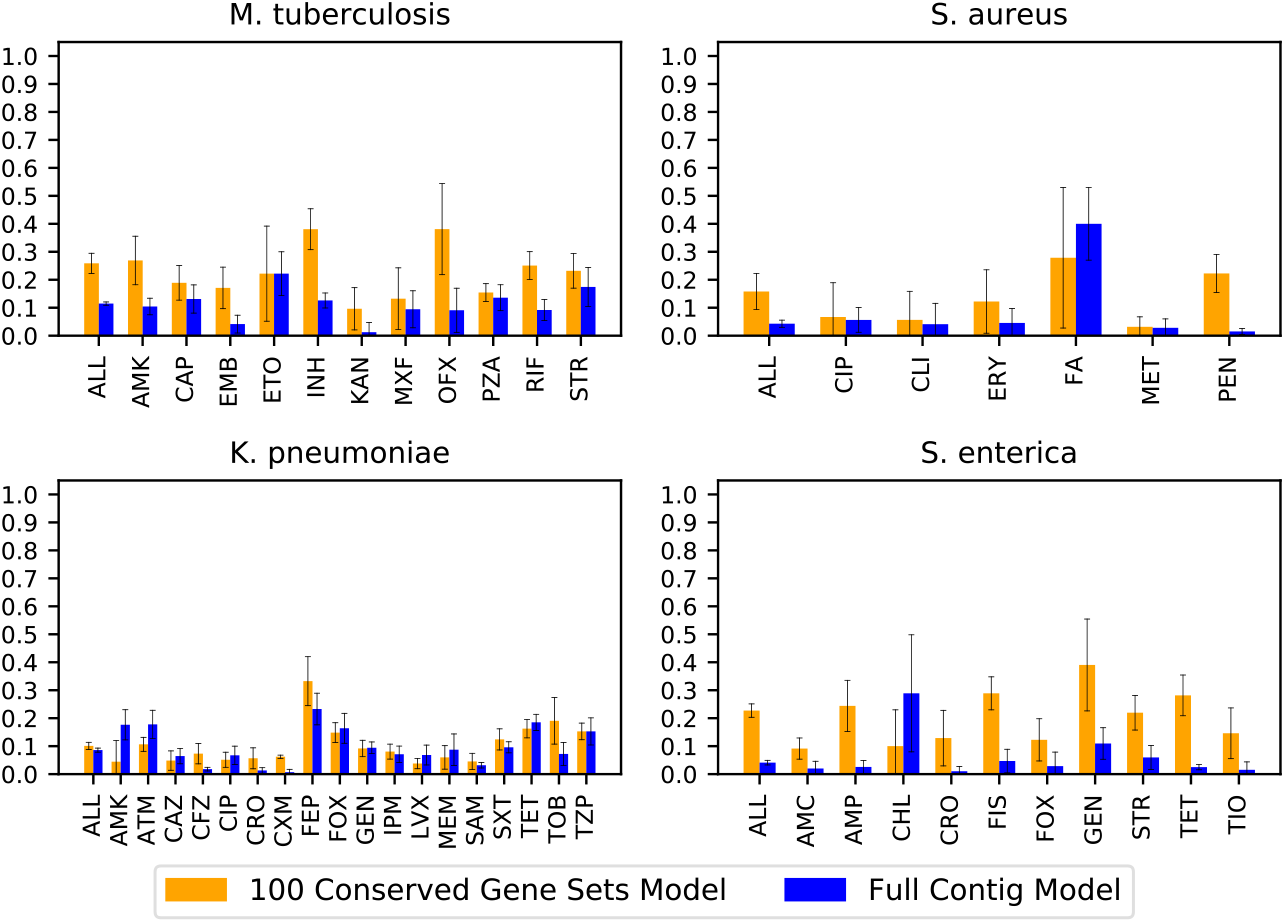
Average very major error rate (VME) by antibiotic for ten models built from non-overlapping sets of 100 randomly-selected core genes (orange bars) compared with the VME for models built from whole assembled contigs for the same genomes (blue bars). VMEs are defined as resistant genomes being incorrectly classified as susceptible. Error bars represent 95% confidence intervals for the whole genome model, and the minimum and maximum observed in all ten replicates for the core gene models.

**Figure 4.**
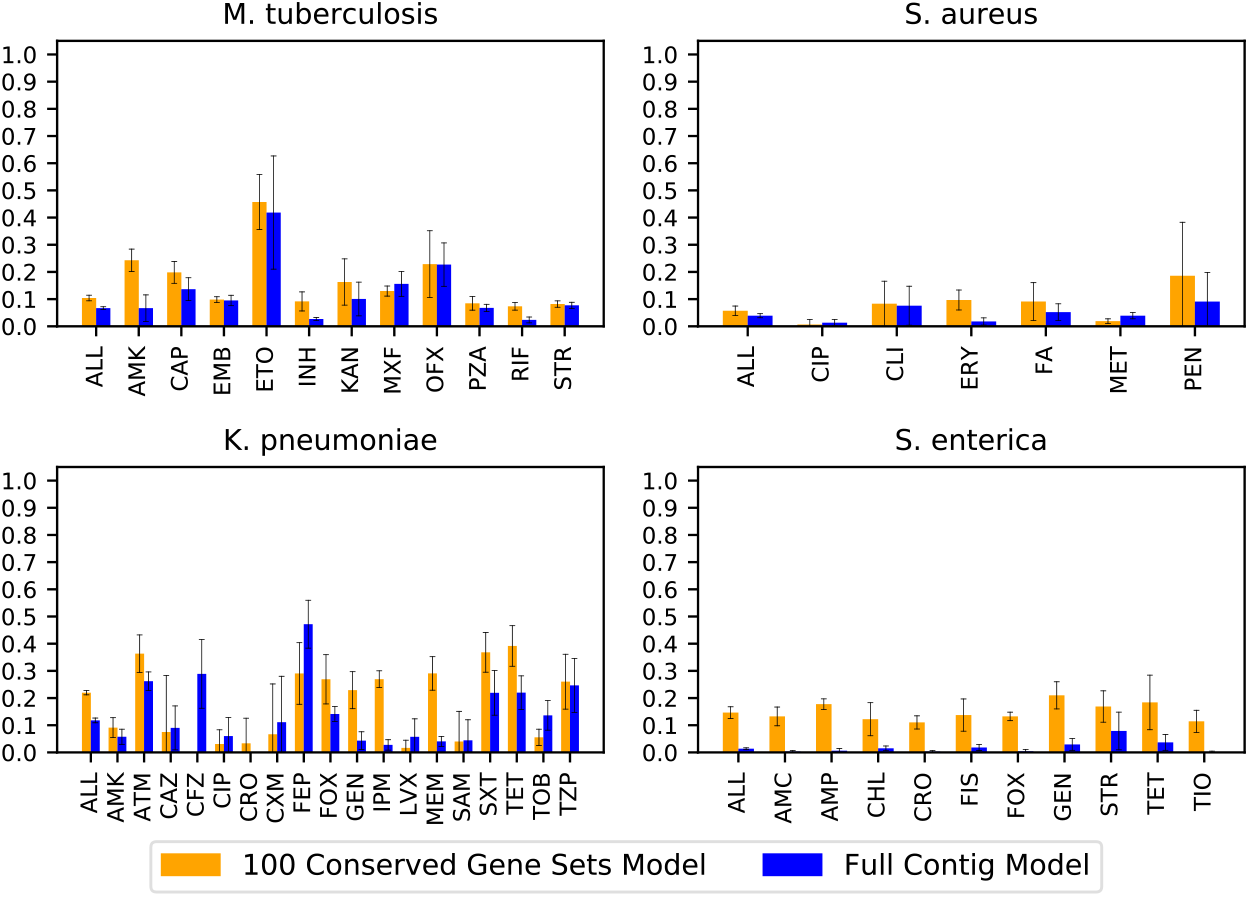
Average major error rate (ME) by antibiotic for ten models built from non-overlapping sets of 100 randomly-selected core genes (orange bars) compared with the ME for models built from whole assembled contigs (blue bars). MEs are defined as susceptible genomes being incorrectly classified as resistant. Error bars represent 95% confidence intervals for the whole genome model, and the minimum and maximum observed in all ten replicates for the core gene models.

### Core gene models are learning nucleotide variation

Although we monitored all models using a validation set to prevent overfitting, it is possible that some aspect of the algorithm or approach, rather than the underlying nucleotide sequences, could be causing the high accuracies observed in the core gene set models. One possibility is that the models are capable of memorizing the data set, rather than learning the nucleotide variation associated with AMR phenotypes. If this were true, we would observe high accuracies regardless of how the genomes are labeled. To test this, we built ten models for ten randomly selected non-overlapping sets of 100 core genes as described above. We then shuffled the labels (i.e., the phenotypes) prior to training the models, and measured the resulting F1 scores (Figure 5). The average F1 scores for the models built from shuffled labels falls to approximately 50%, which is what would be expected from a random guess. This indicates that sequences associated with each phenotype are important for generating an accurate prediction and that the model is not memorizing the data set.

**Figure 5.**
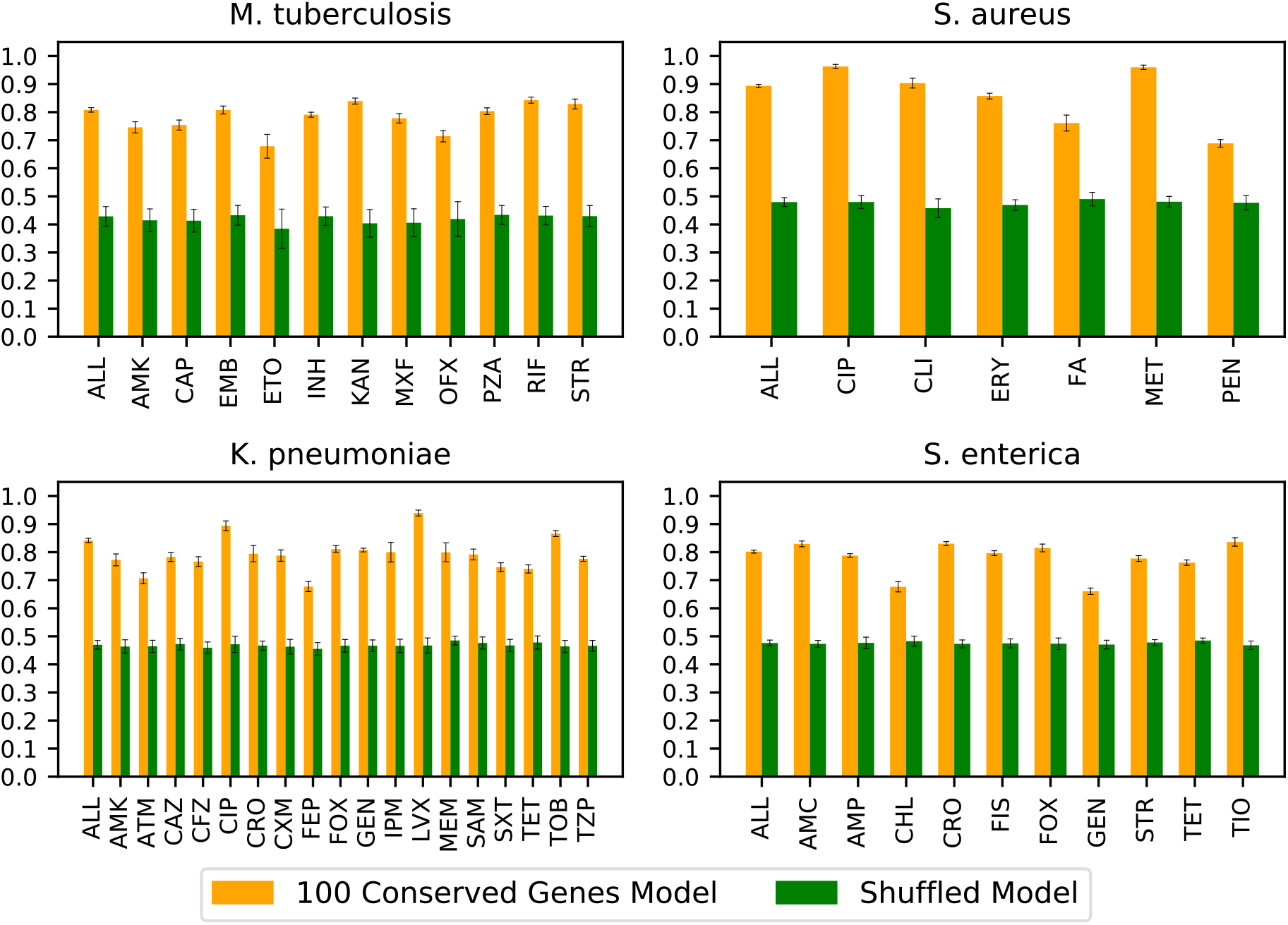
Average F1 scores by antibiotic for ten models built from non-overlapping sets of 100 core genes (orange bars) versus the same model where the AMR phenotypes have been randomized (green bars). Error bars represent the minimum and maximum 95% confidence observed in all ten replicates.

In order to see how the average nucleotide length of a core gene set influences the accuracy of the models, we sorted all of the core genes by length and then built models from 100 genes with average lengths increasing size from 175 to 2,000 nucleotides (Figure S2). For each species except *S. aureus*, we observe a modest increase in F1 scores as the average nucleotide length of the core gene set increases from 175 to 2,000 nucleotides. *K. pneumoniae* had the largest improvement in F1 scores, going from 0.80 [0.79-0.81, 95%CI] for the set averaging 175 nucleotides in length to 0.87 [0.87-0.88, 95%CI] for the set averaging 2000 nucleotides in length. We also observe modest corresponding decreases in the error rates as the length increases. Overall, this suggests that the longer genes may contain more sequence variation that can be found by the machine learning algorithm. However, since the overall gain in accuracy is modest, it also implies that certain core genes may contain more usable variation than others, regardless of their lengths.

### Experimental approach has little effect on core gene models

It is possible that some unforeseen aspect of the experimental design, such as using k-mers as features or XGB as the machine learning algorithm, could be resulting in the high accuracies observed for the core gene models. To control for this, we built a concatenated alignment of 100 randomly-selected core genes for each species. The alignment was one-hot encoded (i.e., converted to a binary vector) and used to build a matrix with the one-hot encoded phenotypes for each species. Models were trained using both the XGB and Random Forest (RF) algorithms. The resulting F1 scores and accuracies for the alignment-based models built using XGB and RF are nearly identical, with overlapping 95% confidence intervals (Figure 6). The F1 scores from the k-mer-based and alignment-based XGB models built from the same set of genes are also nearly identical (Figure 6). The strong agreement between the XGB models built using k-mers or alignment columns as features indicates that feature selection is not biasing the outcome, and that both approaches are using the underlying nucleotide sequence to make the prediction. Furthermore, the close agreement between the XGB and RF models indicates that the high accuracies are not an anomaly relating the algorithm choice, with the slight variation in F1 scores, ME, and VME rates being more consistent with what would be expected from comparing results from different machine learning algorithms.

**Figure 6.**
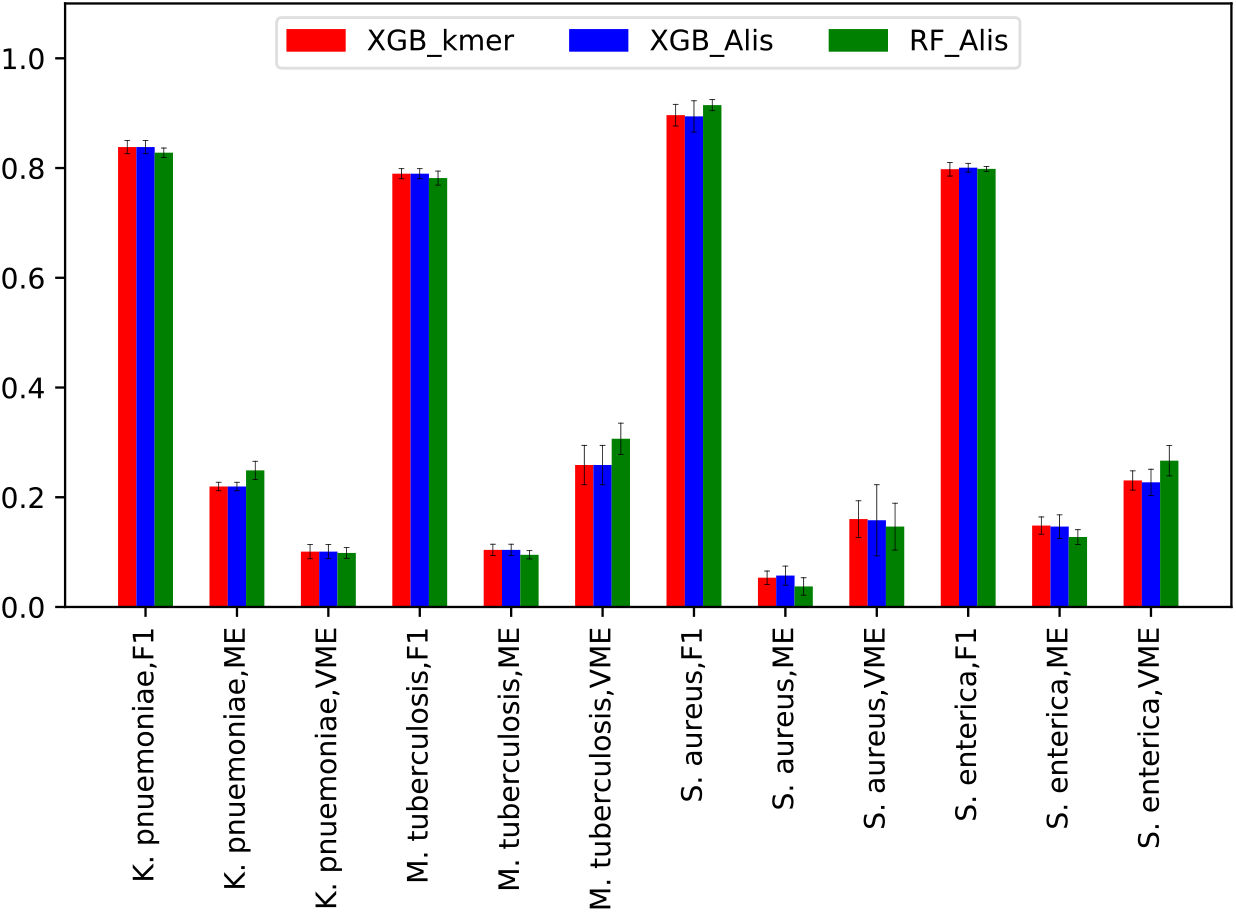
A comparison of algorithms and features. The same set of 100 randomly-selected core genes was used to build each model. Red bars depict the F1 scores for XGBoost models built from k-mers for the gene set. Blue bars depict the F1 scores for a XGBoost model built using a concatenated alignment of the core gene set, where the alignment columns were one hot encoded. Green bars depict the F1 scores for a random forest model using a concatenated alignment of the gene set, where the alignment columns were one hot encoded. Error bars depict the 95% confidence interval.

One aspect of the experimental design that could be increasing the reported accuracy of the core gene models is our decision to build a single model for predicting all AMR phenotypes, rather than building an individual model for each antibiotic. In previous work, we observed that combining antibiotics may result in slightly more accurate models, presumably because related antibiotics lend support to each other in the combined model^22^. To evaluate this, we built individual models for each antibiotic using the same set of 100 randomly selected genes that was used in Figure 1 (Figures S3-5). The F1 scores averaged over all individual antibiotic models is 2-8% lower than the average F1 scores for the merged models. However, when the 95% confidence intervals are compared for each antibiotic, they mostly overlap between the merged and individual models. Overall, combining all antibiotics into a single model slightly improves the F1 scores in some cases, but this is not sufficient to explain why the small non-AMR core gene models have accuracies exceeding 80%.

In previous work, we established methods for predicting minimum inhibitory concentrations (MICs) from whole genome sequence data^21,22^. Although we lack MIC data for *M. tuberculosis* and S. aureus, we also built MIC models using a randomly selected subset of 100 core genes for *K. pneumoniae* and *S. enterica*. In the case of *K. pneumoniae*, the average accuracy of the model within ± 1 two-fold dilution step is 0.87 [0.86-0.87, 95%CI], and is 5% lower than the corresponding whole genome model 0.92 [0.91-0.92, 95%CI] (Table S9). In the case of *S. enterica,* the core gene MIC prediction model has an accuracy of 0.74 [0.73-0.74, 95%CI], and is 17% lower than the accuracy the corresponding whole genome model, which is 0.91 [0.90-0.93, 95%CI]. For both species, the corresponding error rates are also higher for the MIC models built from 100 core genes (Table S9). The 5% drop in accuracy between the whole genome and core gene MIC models for *K. pneumoniae*, and the 17% drop in accuracy for *S. enterica,* are both consistent with the difference in accuracies observed for the SR models described in Table S7, which are 3% and 15% lower for *Klebsiella* and *Salmonella*, respectively (F1 scores are shown in Figure 2). Thus, the data requirements for predicting MICs and susceptible and resistant phenotypes appear to be similar in these cases. The *Salmonella* data set differs from the other data sets in this study because the susceptible genomes are slightly over represented. Also, in order to compute the models more efficiently, the *Salmonella* data set was down selected from a collection of 5278 genomes to a set of 1999 genomes that was maximally diverse based on their k-mer distances^22^. A combination of these factors, or some other aspect of *Salmonella* biology, may be contributing to the larger drop in accuracy going from the whole genome models to the core gene models.

### The models are making predictions from conserved SNPs

The machine learning models described in this study work by using decision trees to make the AMR phenotype predictions. Each node in a decision tree represents a feature that contributes to the overall prediction for a given genome. It is difficult to know how well a model copes with using highly conserved SNPs that are abundant versus strain specific SNPs that occur infrequently in the population. A model that makes its predictions from infrequently occurring SNPs may be biased toward outlier data and not generalizable. On the other hand, a model that is able to make predictions based on more conserved SNPs may be more generalizable.

To evaluate this, we used the alignments of 100 core genes described in the previous section. Passing over each column in the alignment, we asked if the column contains a SNP, and if so, how frequently each nucleotide occurred in that position. We then eliminated the SNP containing columns, starting with those that contained nucleotides occurring most infrequently, reasoning that the infrequently occurring nucleotides represented strain-specific SNPs. This was then repeated eliminating alignment columns containing nucleotides that occurred fewer than 10, 25, 50, etc. times, with the F1 scores, ME, and VME rates were recorded for each run (Figure 7). For each species, the average F1 scores slightly decrease, and the error rates slightly increase as more SNP-containing columns are removed. However, since this trend is so gradual, we would conclude that SNPs that are conserved still contain a large amount of predictive power, and strain-specific SNPs are not required for making good predictions.

**Figure 7.**
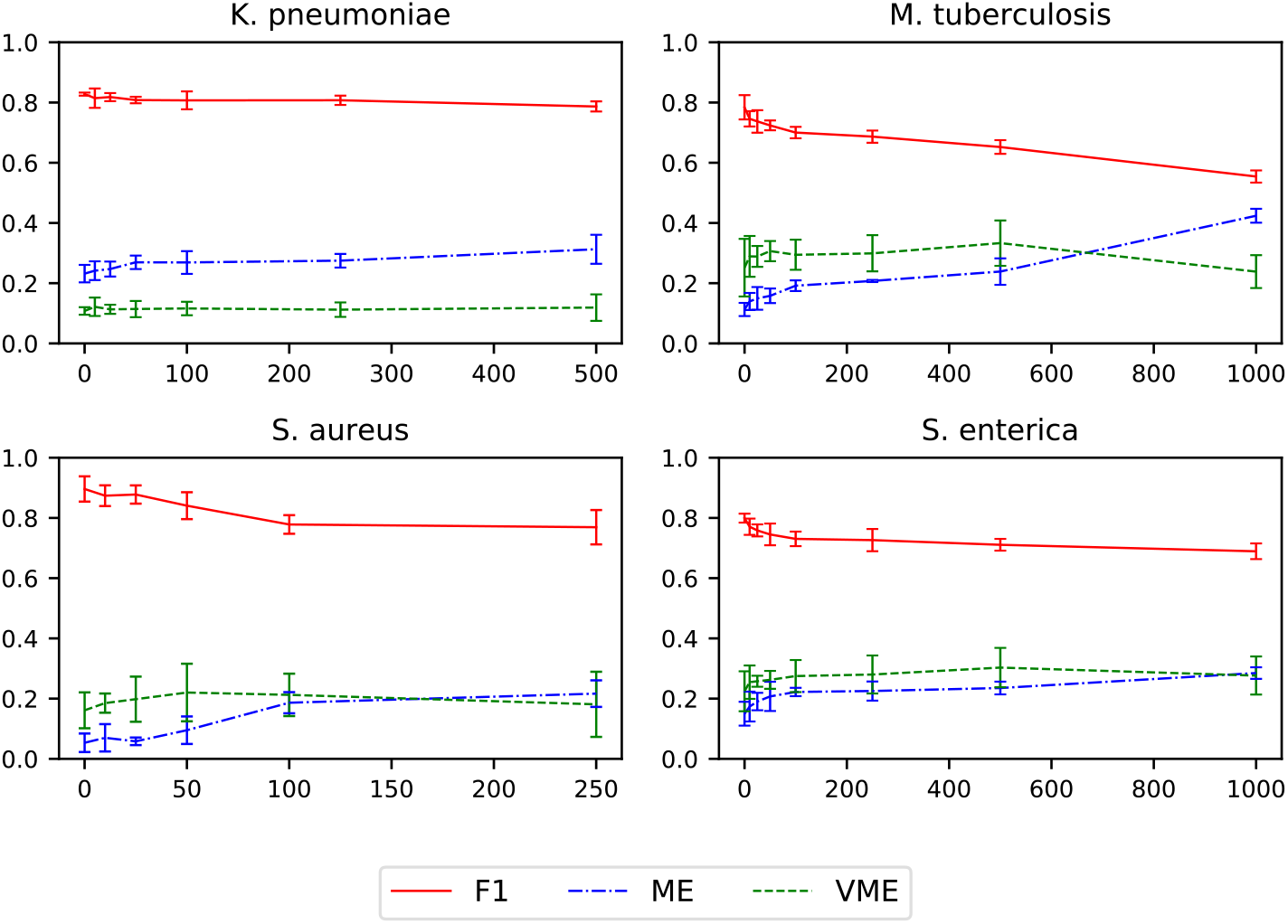
The effect of removing SNPs from core gene models. XGBoost models were built for a randomly selected set of 100 core genes for each species (the same set that was used in Figure 6), using the one-hot encoded concatenated alignment as input. SNP-containing columns with less than 10, 25, 50 etc. nucleotides of the same type were successively removed and a model was generated. The X-axis depicts the SNP conservation and the Y-axis depicts the F1 score. The F1 score (red line), ME rate (blue line), and VME rate (green line) are shown. Error bars depict the 95% confidence interval.

### Strain diversity and phenotype distribution have little influence on the consistency of the models

Strain diversity or sampling may play a role in influencing the accuracy of the models. In the context of a phylogenetic tree, this could happen if many strains are closely related, resulting some clades being much larger than others, or if there is an uneven distribution of susceptible and resistant strains within clades. When this occurs, it is possible that larger clades or susceptible and resistant distributions within clades could bias the models.

To assess how strain diversity influences the models, we computed phylogenetic trees for each species using core genes that were not used in any of the AMR models. We defined clades of varying sizes based on tree distance, and generated k-mer based models, weighting the influence of each genome in the model based on 1) how many genomes are in a given clade, and 2) how many susceptible and resistant genomes occur in a given clade. The weighting was designed to reduce the influence of overrepresented clades or phenotypes in the model. Models were computed using each weighting scheme and a combination of both, using a random sample of 100 core genes.

Overall, for each of the four species, there are no observable differences between the unweighted models and the models that were built from genomes that were weighted according to clade size at various tree distances (Table S10). Similarly, in most cases the 95% confidence intervals overlap for unweighted models and models that were weighted according to the distribution of susceptible and resistant genomes within each clade at each tree distance (Table S10). Combining both weighting schemes also has a nearly identical outcome to weighting according to the distribution of S and R genomes within clades.

Since training and testing at the same tree distance could potentially mask biases, each model that was trained at a given tree distance was also used to evaluate the test set of genomes defined at every other tree distance for a given species, and the F1 scores were averaged by clade (Table S11). For clade-size weighted models, the minimum and maximum 95% confidence intervals for the F1 scores observed for all comparisons in this analysis were 0.79-0.89 for *K. pneumoniae,* 0.78-0.91 for *M. tuberculosis*, 0.75-0.86 for *S. enterica*, and 0.84-0.96 for *S. aureus.* SR weighting had a similar outcome, with the minimum and maximum 95% confidence intervals for any comparison being 0.74-0.87 for *K. pneumoniae*, 0.72-0.90 for *M. tuberculosis*, 0.75-0.86 for *S. enterica*, and 0.83-0.94 for *S. aureus.* The range in F1 scores indicates that there is variation between clades, with the phenotypes of the genomes in some clades being easier to predict that than others. However, the results do not indicate extreme drops in accuracy or imbalances that would indicate that the input data are biasing the model based on clade size or S and R phenotype distributions. In other words, the ability to predict S and R phenotypes is relatively consistent over the tree.

### Feature importance for core gene sets

Because of the relatively high accuracy of the AMR models built from small sets of core genes, and their consistency across random samples, we wanted to see if the models were relying on genes occurring in similar chromosomal regions, or on genes with related functions. To do this, we used the ten models that were built from 100 randomly sampled non-overlapping core gene sets described in Figures 2-4. Each k-mer used in a model has an associated feature importance value that describes its contribution to the model. The feature importance for the k-mers of each gene were summed to approximate the relative contribution of each gene to each model.

We plotted each gene based on its chromosomal coordinates in a reference strain, coloring each gene by its total feature importance (Figure 8). The core genes tend to distribute evenly over the entire chromosome, but there are several regions that lack important genes. These regions will contain non-coding regions, such as the ribosomal RNA operons, and unconserved regions, such as prophage, which would not have been sampled. Overall, for any individual model, we do not observe clustering of important genes on the chromosome, and we also observe no clustering of important genes across models.

**Figure 8.**
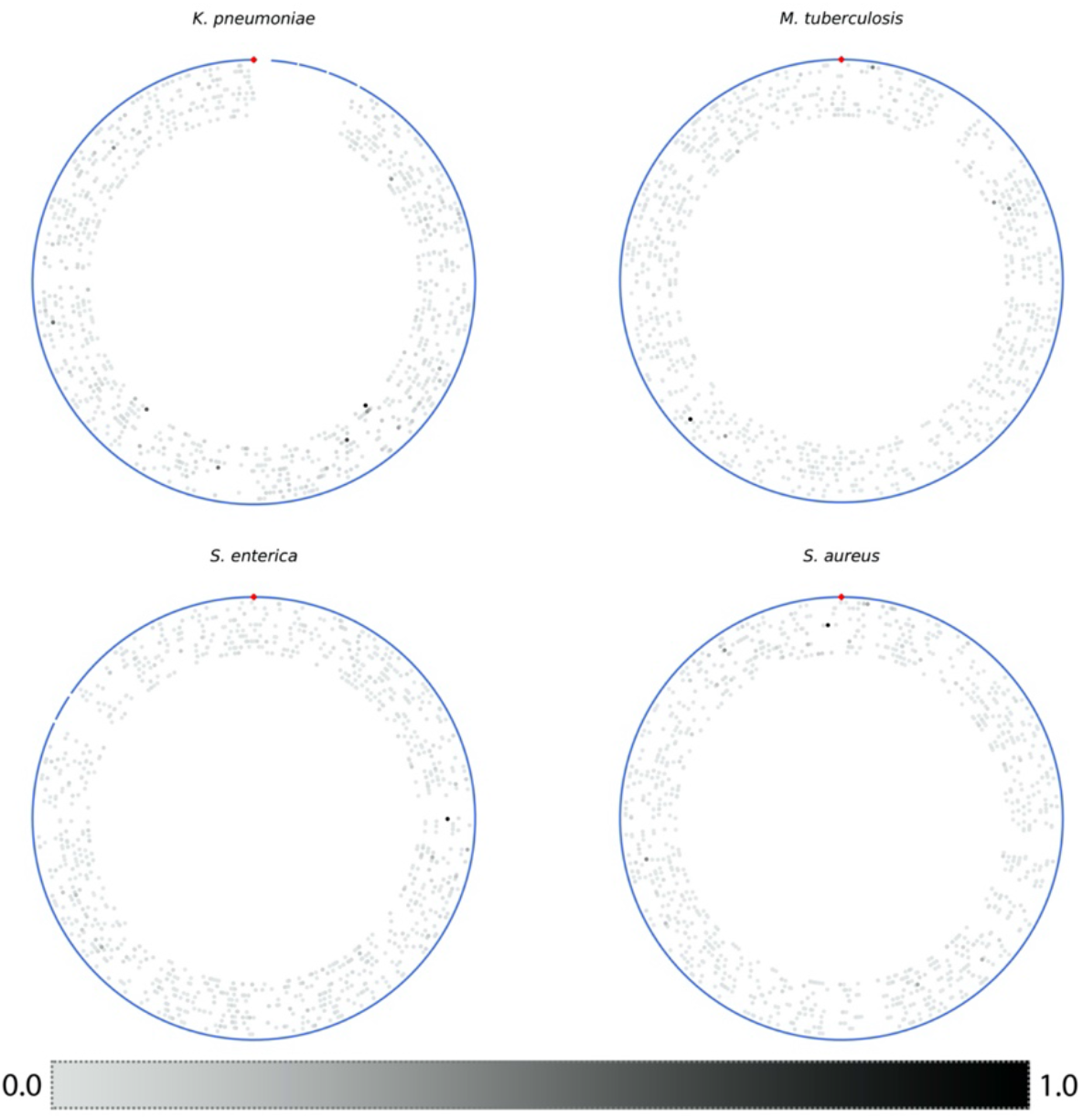
Plots showing the position and total feature importance of each core gene based on its coordinates in a reference genome. The contigs from the reference genome are depicted as blue lines on the outside of each plot, and the origin of replication is shown at the top with a red diamond. Each gene is depicted as a gray dot, and the genes with higher total feature importance in each model are colored in darker shades of gray. The genes of each model are shown as concentric circles from outside to inside. The reference strains used were *K. pneumoniae* HS11286, *M. tuberculosis* H37Rv, *S. enterica* serovar Typhimurium LT2, and *S. aureus* NCTC 8325.

For each set of 100 genes, the relative contribution of each k-mer, and thus each gene, to the model is not uniform. A small number of genes tend to contribute most of the predictive k-mers to each model. This is followed by a relative decline in feature importance over the set (Figure S6), and in some sets, there are genes that do not have a k-mer that is used in the model (Table S8). In several cases, these genes with low or non-contributing k-mers include ribosomal protein genes, which tend to have lower total feature importance scores in the models, presumably due to their slower mutation rates. The ensemble machine learning methods can be frugal in selecting important features, so these feature lists may not include a comprehensive set of k-mers that could have provided signal. Regardless, the highly-ranking k-mers appear to indicate the presence of genes with important biological properties that correlate with the presence or absence of AMR in the population.

For each species, the three genes with the highest total feature importance are shown in Table 3. Most of these genes do not have an obvious role in AMR (Table 3, Table S8), although this is not surprising because the well annotated AMR genes were discarded in this study. Ascertaining if these genes have a role in AMR is challenging, but some clues can be gleaned from web resources and the literature. For example, we used the MycoBrowser^50^ web resource to examine the top gene from each of the ten core gene models for *M. tuberculosis.* Nine of the top ten genes were listed as non-essential in laboratory transposon mutagenesis studies^51–53^, which is surprising given their strong conservation in over 5,000 strains. One *M. tuberculosis* gene, OpcA (Rv1446c), was listed as essential^52,54^ and was found to be upregulated in isoniazid resistant clinical isolates in a proteomic study^55^, which may imply a role in AMR. Six of the of the top ten *M. tuberculosis* genes also encode proteins that are associated with the cell membrane or wall^56^, which could imply roles in biofilm formation, immune system avoidance, transport, or virulence. This is also true for several of the other genes listed in Table 3. For example, the *wzi* gene (annotated as 55.8kDa ORF3) in *K. pneumoniae* is involved in the K-antigen capsular polysaccharide formation, and is an important virulence factor^57–60^; the staphylococcal secretory antigen *ssaA*, is involved in biofilm formation^61,62^ and is also a likely virulence factor in Staphylococcus strains^63^; and the large *Salmonella bapA* gene (annotated as T1SS secreted agglutinin RTX) is also a virulence factor that is necessary for biofilm formation^64^. Although perhaps not an AMR gene *per se*, the *S. aureus rlmH* gene (annotated as 23S rRNA (pseudouridine(1915)-N(3))-methyltransferase (EC 2.1.1.177)), contains the integration site for SCC*mec* elements, which confer methicillin resistance^48,65,66^. Indeed, the highest-ranking k-mer for this gene, “AAGCATATCATAAAT”, corresponds with the integration site and has a feature importance score of 389, which is 80% of the total feature importance for this gene. Upon integration, the attachment site, *attB*, is split into two similar sequences, *attL* and *attR*^65^, and this k-mer is picking up this key difference between susceptible and resistant strains.

**Table 3.** The top three genes with the highest total feature importance from all k-mers used in each model. Results are based on ten separate models built from 100 randomly sampled non-overlapping core gene sets.

| Protein family | Average gene length | Cumulative feature importance | PATRIC annotation |
| --- | --- | --- | --- |
| ***K. pneumoniae*** |  |  |  |
| PLF_570_00002496 | 1422.3 | 818.8 | Uncharacterized 55.8 kDa protein in cps region (ORF3) |
| PLF_570_00001044 | 1419.6 | 669.2 | Alpha,alpha-trehalose-phosphate synthase [UDP-forming] (EC 2.4.1.15) |
| PLF_570_00003328 | 1247.9 | 533.3 | N-carbamoyl-L-amino acid hydrolase (EC 3.5.1.87) |
| ***M. tuberculosis*** |  |  |  |
| PLF_1763_00001229 | 909.9 | 1157.0 | OpcA, an allosteric effector of glucose-6-phosphate dehydrogenase, actinobacterial |
| PLF_1763_00001419 | 697.0 | 649.9 | putative phosphoglycerate mutase |
| PLF_1763_00001681 | 1580.9 | 518.2 | DNA methylase |
| ***S. enterica*** |  |  |  |
| PLF_590_00006292 | 11345.6 | 1828.8 | T1SS secreted agglutinin RTX |
| PLF_590_00001168 | 1593.0 | 616.6 | Bis-ABC ATPase YbiT |
| PLF_590_00001455 | 906.0 | 571.3 | GTP-binding protein Era |
| ***S. aureus*** |  |  |  |
| PLF_1279_00000411 | 477.3 | 484.8 | 23S rRNA (pseudouridine(1915)-N(3))-methyltransferase (EC 2.1.1.177) |
| PLF_1279_00002023 | 687.0 | 250.0 | Phage-encoded chromosome degrading nuclease YokF |
| PLF_1279_00001560 | 899.8 | 232.0 | Secretory antigen precursor SsaA |

## Discussion

In this work, we have shown that AMR phenotypes can be predicted from sets of core genes that are not annotated to be involved in AMR. These gene sets can be quite small, less than 100 genes, and still have predictive power. Building models from randomly sampled non-overlapping sets of 100 core genes results in model accuracies that are quite stable with little variation between samples. This result is not influenced by the choice of features (nucleotide k-mers or alignments) or machine learning algorithms (XGB or RF), and the high accuracies do not appear to be the result of overfitting, memorization, strain-specific SNPs, or imbalances in sampling or phylogeny. This effect is seen in the Gram-negative *K. pneumoniae* and *S. enterica* genomes, and the Gram-positive *M. tuberculosis* and *S. aureus* genomes, indicating that this is likely widespread. These small core gene models also have predictive power in the cases where the primary AMR mechanism results from SNPs or horizontal gene transfer.

Although the small core gene models described in this study do not meet the accuracy or error rate requirements of clinical diagnostic tools^39^, the results have implications for the development of bioinformatic strategies aimed at tracking and understanding AMR. The models demonstrate that the prediction of AMR phenotypes from incomplete genome sequence data is possible, even when the AMR genes are missing. This means that AMR phenotype predictions could be made for contigs that are binned from a metagenomic assembly, even when they do not represent a complete chromosome, or lack the corresponding plasmid contigs that carry important AMR genes. This assertion comes with the obvious caveat that using more sequence data in the training set results in models with higher accuracies and lower error rates. In order to demonstrate the predictive power of these small core gene models, we specifically chose to avoid using well-known AMR genes, and we focused on smaller gene sets. However, it is possible to envision a variety of approaches that could improve the accuracy of the models or result in more rapid AMR prediction. For instance, accuracies might be improved by training on larger core gene sets, training on select chromosomal regions, or by finding bespoke conserved gene sets that optimize the predictive power. Predictions could be made more rapid by employing read mapping strategies against specific gene sets to avoid de novo assembly.

Although this study was not designed to be an exhaustive search for uncharacterized AMR genes, it does highlight the value of the large publicly-available genomic data sets that have paired AST metadata, and the power of using AI methods for identifying important information within them. In particular, the analysis of important features in the core gene models revealed at least two genes, *rlmH* and *optA*, with a known or possible role in AMR. Several of the other genes that were identified as contributing important k-mers were well-known virulence factors including *wzi*, *bapA*, and *ssaA*. It is difficult to know if the genes encoding these virulence factors have a direct biological role in AMR, or if certain alleles of the gene simply correlate with resistant or susceptible genomes. In other words, the resistant genomes might also be expected to be the more virulent genomes and vice versa.

Another consideration is the potential of the core gene models for identifying compensatory or epistatic changes throughout the genome. These mutations occur in non-AMR genes in order to accommodate the potentially reduced fitness cost of maintaining the primary AMR conferring genes or SNPs^67–70^. The results of this study, and previous AI studies focusing on AMR^25,27^, suggest that these changes could be widespread throughout the genome. Indeed, although each model had a few genes with very high total feature importance values, most genes were contributing k-mers to the decision trees for each model. In general, AMR-related epistatic changes remain poorly understood, and are perhaps combinatoric, but an incisive use of AI methods may help to establish a more mechanistic understanding of this phenomenon.

## Abbreviations

AI: artificial intelligence
CI: confidence interval
ME: major error, susceptible genomes predicted to be resistant
MIC: minimum inhibitory concentration
PATRIC: Pathosystems resource integration center
PLF: PATRIC local protein family
RAST: Rapid annotation using subsystem technology
RF: Random Forest
SR: susceptible and resistant
VME: very major error, resistant genomes predicted to be susceptible
XGB: XGBoost

## Acknowledgements

We thank the members of the BV-BRC PATRIC, iSENTRY, NARMS and Houston Methodist teams, Dion Antonopoulos, Philippe Noirot, and Sarah Owens for their helpful comments. We thank NARMS and Houston Methodist for the *Salmonella* and *Klebsiella* data sets. We also thank Emily Dietrich for her careful editing. This work is funded by the United States Defense Advanced Research Projects Agency iSENTRY Friend or Foe program award [Contract No. HR0011937807], and by the United States National Institute of Allergy and Infectious Diseases Bacterial and Viral Bioinformatics resource center award [Contract No. 75N93019C00076].

## Author Contributions

MN: study design, data generation, manuscript preparation; RO: genome assembly; MS: AMR metadata integration, genome annotation, MV: AMR metadata curation, JD study design, data generation, manuscript preparation

## Supplementary Figures

**Figure S1.**
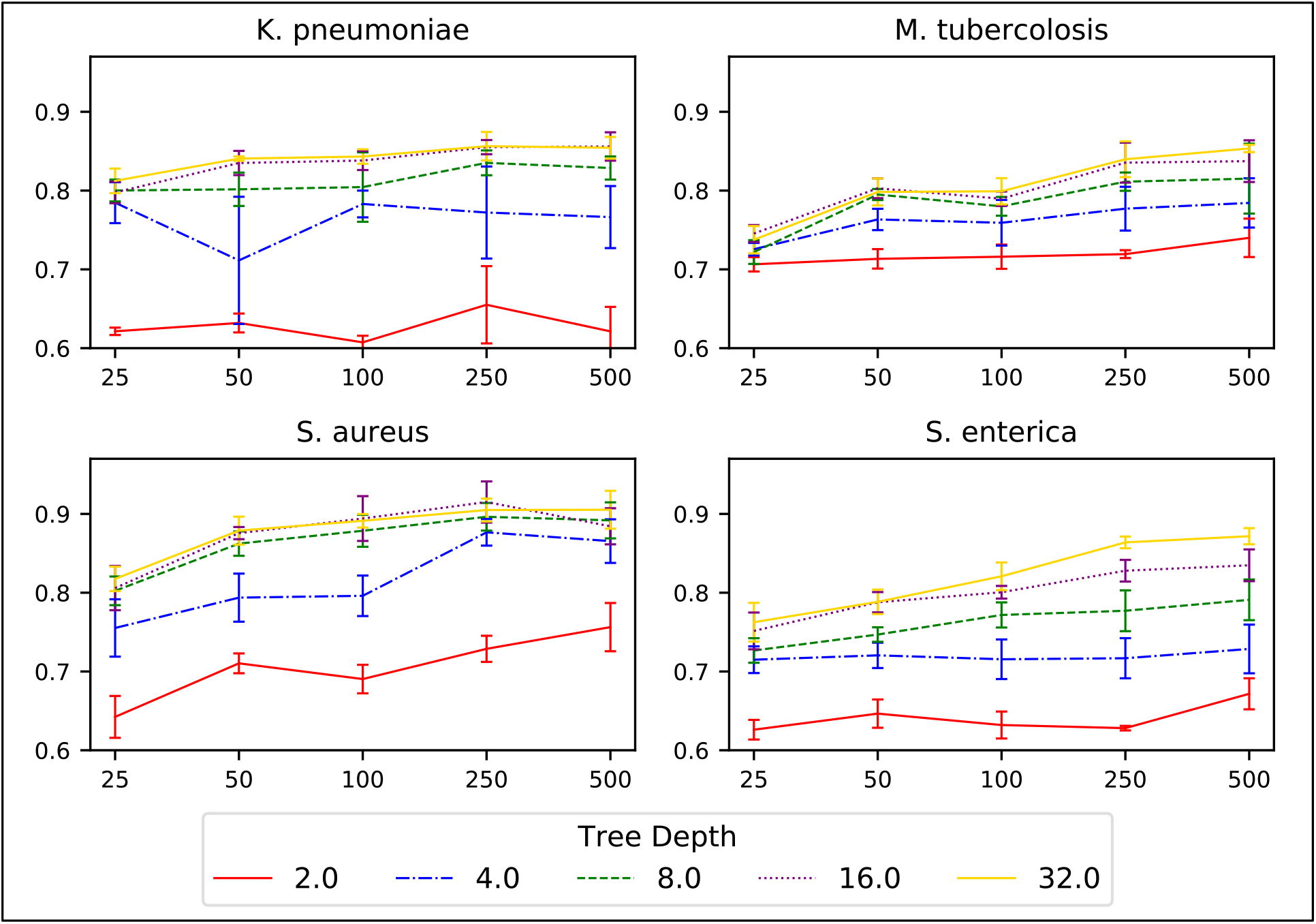
The effect of tree depth on models built from core gene sets of various sizes. K-mer-based XGB models were built for randomly-selected core gene sets ranging in size from 25-500 genes. The X-axis depicts the core gene set size, and the Y-axis depicts F1 scores. Each line represents models built at varying tree depths from 2-32. Error bars are the 95% confidence interval.

**Figure S2.**
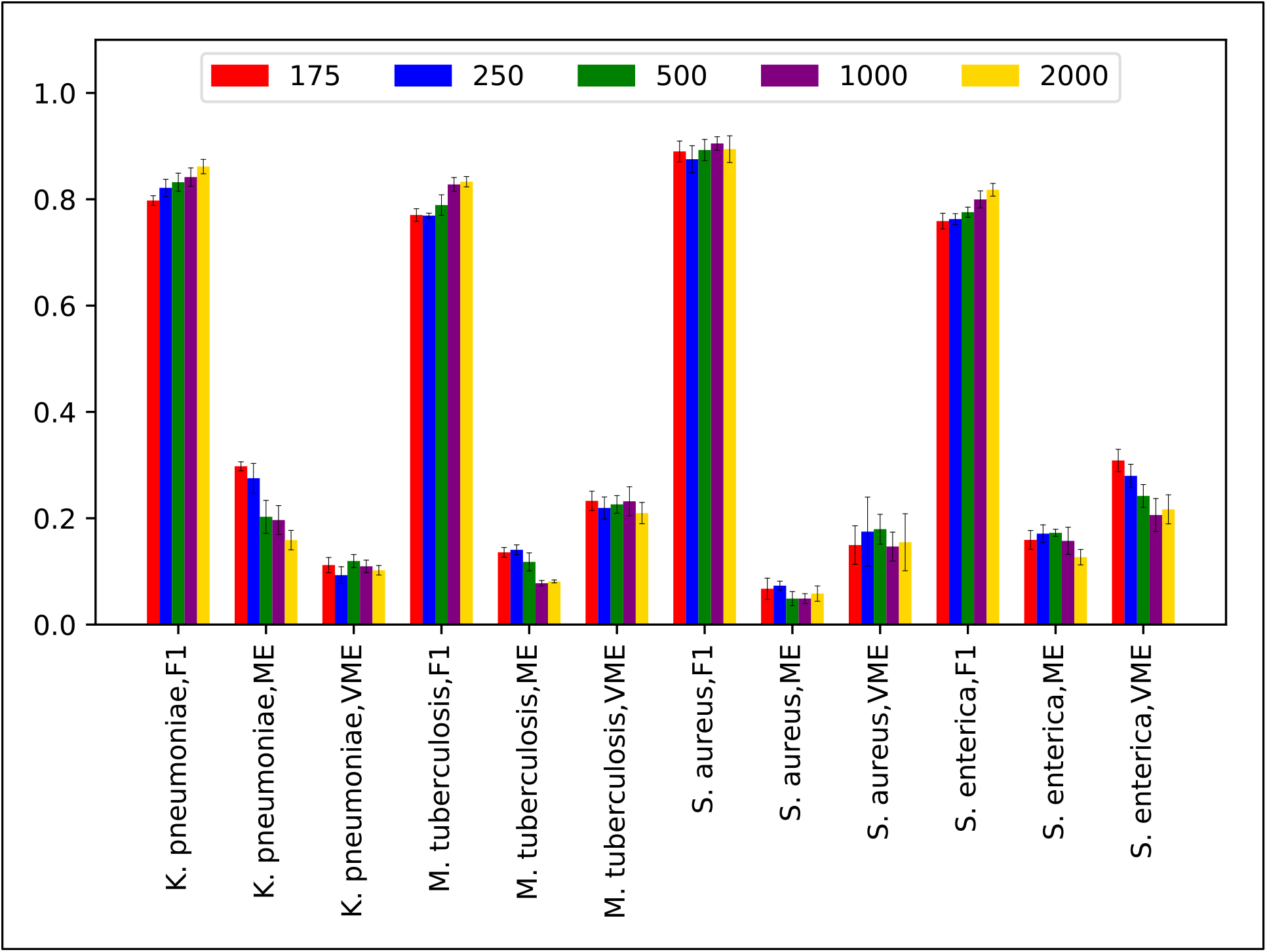
The effect of building models from core gene sets of varying nucleotide lengths. Core genes were sorted by length and k-mer-based XGB models were built for the set of 100 genes with average lengths that were closest to 175, 250, 500, 1,000, and 2,000 nucleotides. Bars show F1 scores, ME rates, and VME rates. Error bars are the 95% confidence intervals.

**Figure S3.**
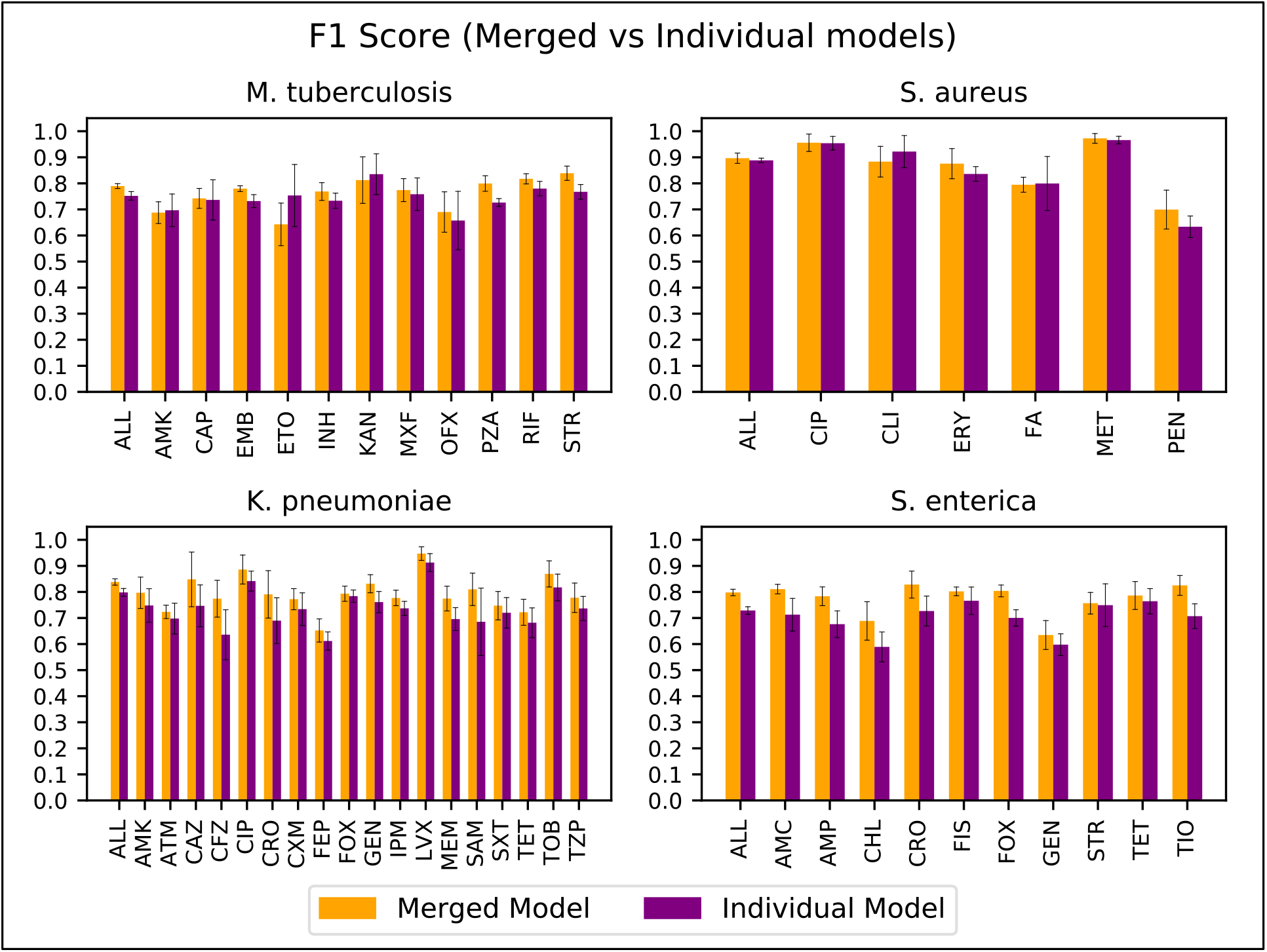
A comparison of F1 scores for merged models, where all antibiotics are included in a single model, and individual models built for each antibiotic. K-mer-based XGB models were built from the same set of 100 randomly selected core genes. Error bars represent the 95% confidence intervals.

**Figure S4.**
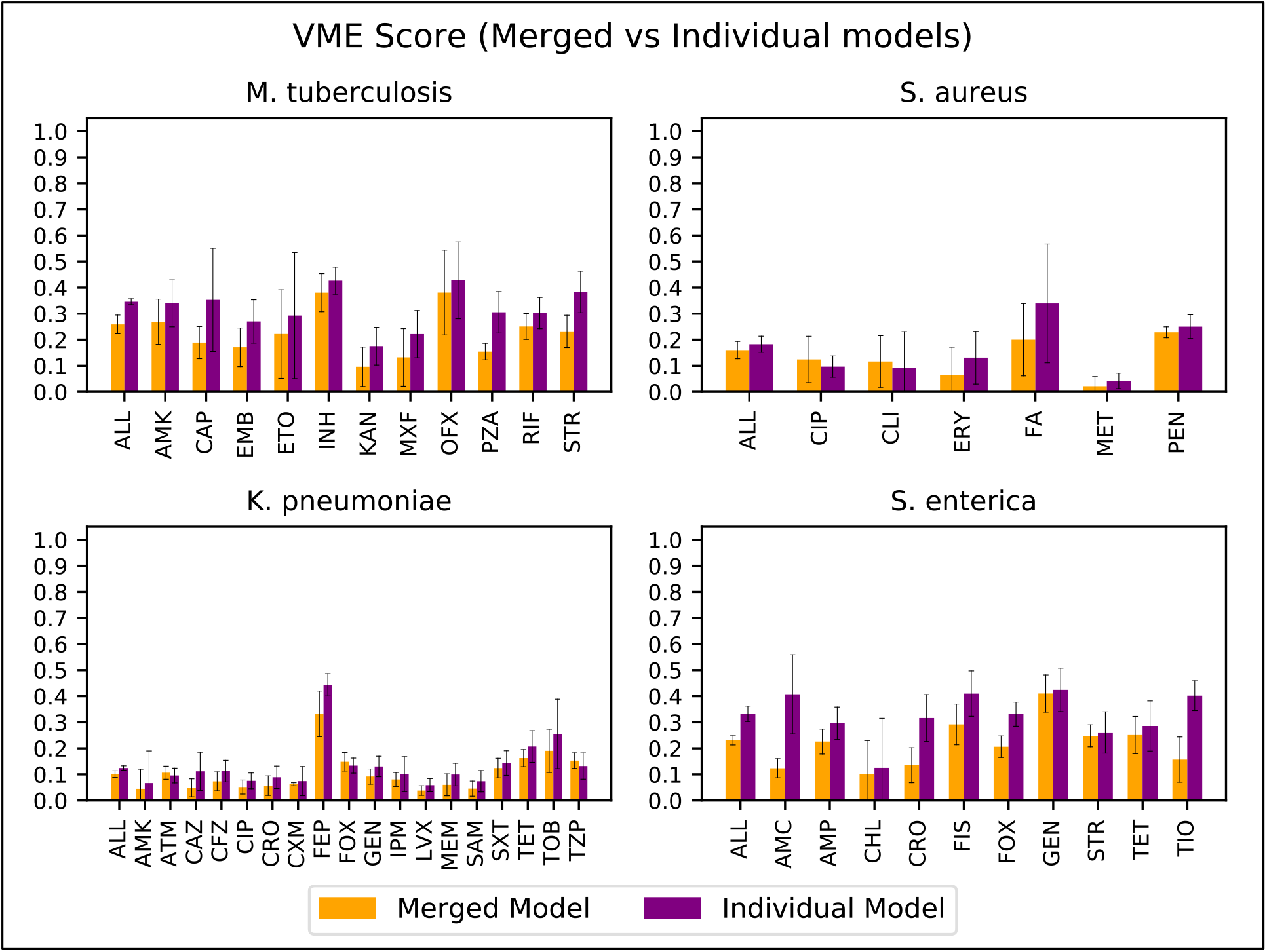
A comparison of very major error rates for merged models, where all antibiotics are included in a single model, and individual models built for each antibiotic. K-mer-based XGB models were built from the same set of 100 randomly selected core genes. Error bars represent the 95% confidence intervals.

**Figure S5.**
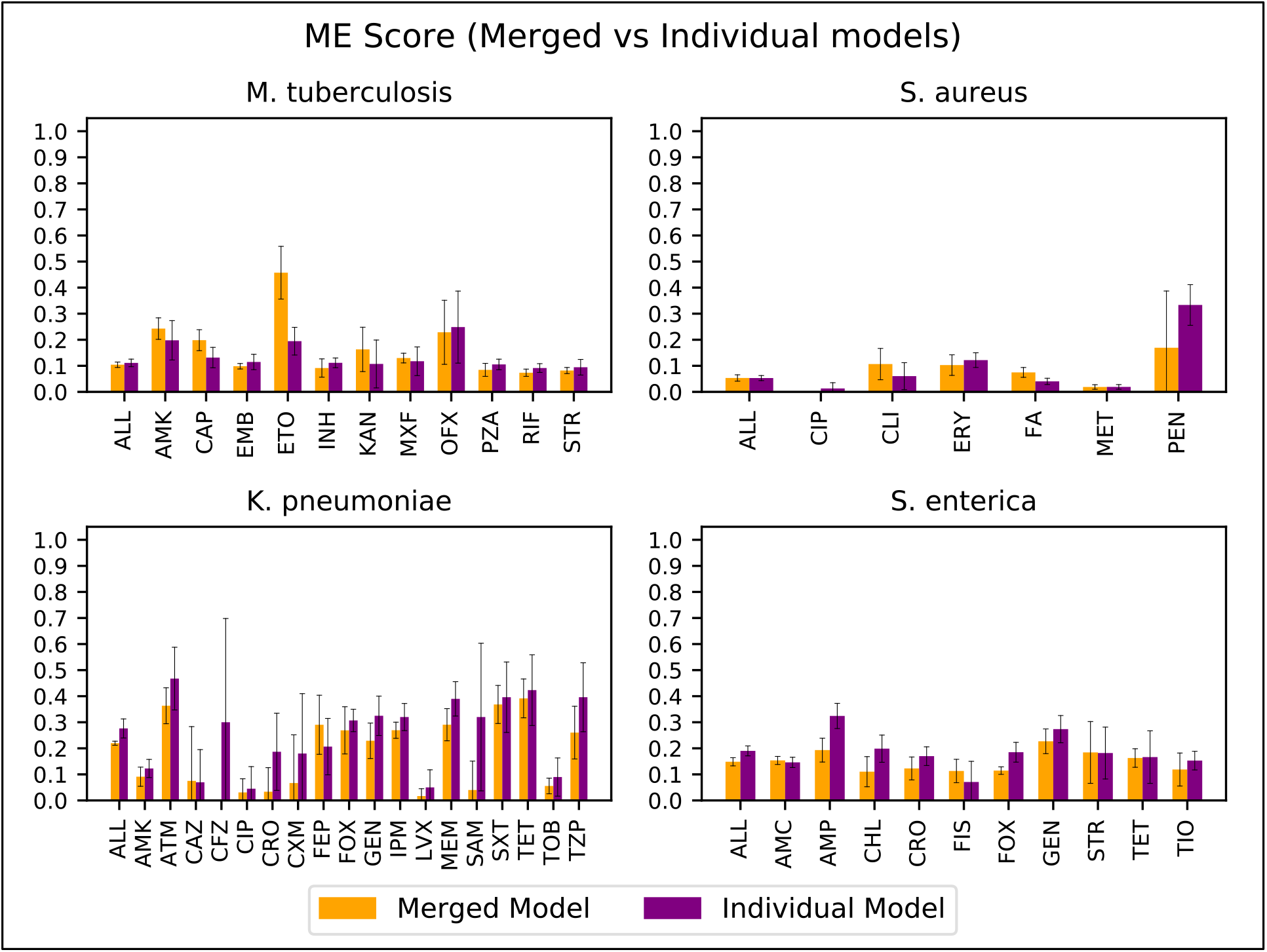
A comparison of major error rates for merged models, where all antibiotics are included in a single model, and individual models built for each antibiotic. K-mer-based XGB models were built from the same set of 100 randomly selected core genes. Error bars represent the 95% confidence intervals.

**Figure S6.**
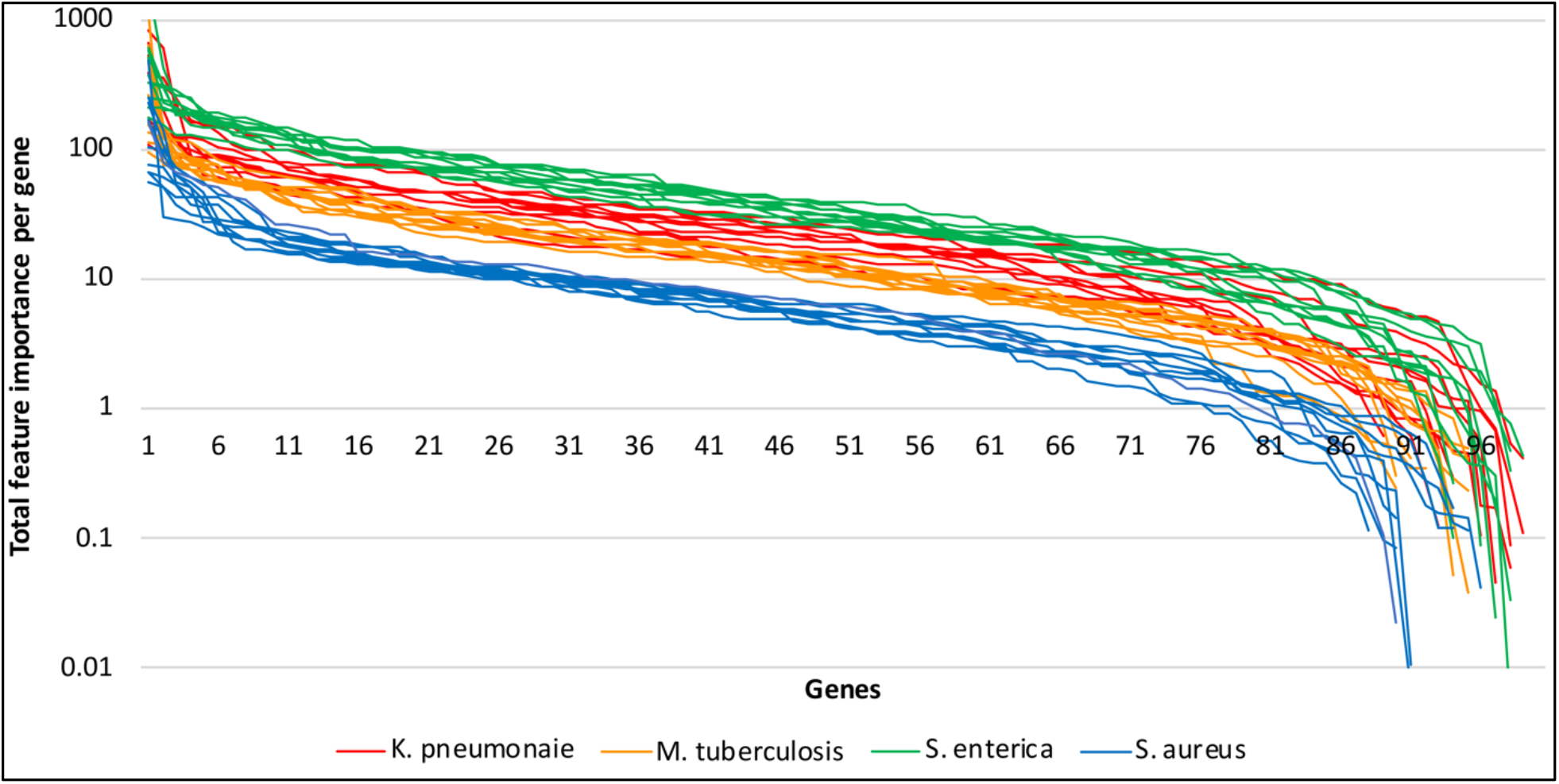
Total feature importance per gene. K-mer-based XGB models were built for 10 non-overlapping sets of 100 core genes. The X-axis depicts each of the 100 genes sorted by total feature importance and the Y-axis is the total feature importance score. Each line depicts one model for *K. pneumoniae* (red), *M. tuberculosis* (orange), *S. enterica* (green), and *S. aureus* (blue).

